# Human endogenous retrovirus-R envelope is a host restriction factor against severe acute respiratory syndrome-coronavirus-2

**DOI:** 10.1101/2022.08.05.502940

**Authors:** Nidhi Gupta, Shabnam Ansari, Rohit Verma, Oinam N Singh, Mukesh Kumar Yadav, Akshay Binayke, Kamini Jakhar, Shailendra Mani, Amit Awasthi, Shalimar, Baibaswata Nayak, C.T. Ranjith-Kumar, Milan Surjit

**Author notes:** Correspondence: Milan Surjit. These authors contributed equally (Nidhi Gupta, Shabnam Ansari). Department of Biochemistry, All India Institute of Medical Sciences, New Delhi, India (Nidhi Gupta).

## Abstract

Coronavirus induced disease-19 (COVID-19), caused by the SARS-CoV-2 remains a major global health challenge. Human endogenous retroviruses (HERVs) represent retroviral elements that got integrated into the ancestral human genome. HERVs are important in development and diseases, including cancer, inflammation and viral infections. Here, we analyzed the expression of several HERVs in SARS-CoV-2 infected cells and observed increased activity of HERV-E, HERV-V, HERV-FRD, HERV-MER34, HERV-W and HERV-KHML2. In contrast, HERV-R-envelope was downregulated in cell-based models and COVID-19 patient PBMCs. HERV-R overexpression inhibited SARS-CoV-2 replication, suggesting its antiviral action. Further studies demonstrated the role of extracellular signal-regulated kinase (ERK) in regulating HERV-R antiviral activity. Cross-talk between the ERK and p38 MAPK controls HERV-R envelope synthesis, which in turn modulates the replication of SARS-CoV-2. These findings establish the importance of HERV-R envelope as a host restriction factor against SARS-CoV-2 and illustrate the advantage of integration and evolutionary maintenance of retroviral-elements in the human genome.

## Introduction

Human genome contains sequences of several ancient retroviruses, which infected and got integrated into the germ line cells during the course of evolution, over millions of years ago. Collectively, these retroviral sequences are categorized as Long-terminal repeats (LTR) retrotransposons. Human endogenous retroviruses (HERVs) account for ∼8% of the human genome (Grandi and Tramontano, 2018a,b; Mayer et al.,2011). HERVs have been classified into three classes and 11 supergroups. Gamma- and Epsilon- retroviruses like HERVs, such as MLLV, HERV-ERI, HERV-FRD, HERV-W etc., belong to the class I. Beta-retrovirus-like HERVs, such as the HERV-K (HML supergroup) belong to class II. Spuma-retrovirus-like HERVs, such as HERV-L and HERV-S (HSERVIII supergroup), belong to class III (Vargiu et al., 2016). Although HERVs were earlier considered “junk DNA sequences”, emerging evidence suggests their significant involvement in normal development and physiological homeostasis as well as multiple diseases (Mao et al., 2021; Grandi and Tramontano, 2018a, b). For example, envelope proteins of HERV-W and HERV-FRD (named as syncytin-1 and syncytin-2, respectively) are expressed in placenta and help in the formation of the syncytiotrophoblast layer of the placenta (Blond et al., 2000; Malassine et al., 2004; Mi et al., 2000). Moreover, different HERVs are transcribed during various stages of embryogenesis, thus acting as hallmarks of embryonic development. Transcriptional activation of HERVs have been observed in many human cancers such as breast cancer, leukemia, hepatocellular carcinoma and neurodegenerative diseases such as multiple sclerosis and type I diabetes (Grandi and Tramontano,2018a).

Many HERVs induce the expression of interferon-stimulated genes (ISGs), thus activating the antiviral innate immune response pathways (Li et al., 2022). HERV-K activates NF-kB, cGAS (cyclic GMP-AMP) and STING (cyclic GMP-AMP receptor stimulator of interferon genes) signaling pathways (Li et al., 2022). HERVs are also implicated in many viral infections, including DNA viruses (such as KSHV, HSV-1, HCMV, EBV), RNA viruses (Hepatitis C Virus, Dengue virus, Influenza virus, SARS-CoV-2) and retroviruses (HIV-1, HTLV) (Dai et al.,2018; Bello-Morales et al., 2021; Assinger et al.,2013; Bergallo et al., 2015; Sutkowski et al., 2001; Tovo et al., 2020; Kitsou et al., 2021; Balestrieri et al., 2021; Bhardwaj et al., 2020; Contreras-Galindo et al., 2006; Wang et al., 2020). Notably, HERV-H- Pol (polymerase) and HERV-K-Pol is overexpressed in chronic HCV infection (Tovo et al., 2020). HERVs are activated in dengue virus infection (Wang et al., 2020). ICP0 protein of HSV-1 upregulates HERV-K (Bello-Morales et al., 2021). EBV infection upregulates the level of the envelope protein of HERV-K18 and HERV-W (Sutkowski et al., 2001; Mameli et al., 2012). HERV-K-HML2 transcription is increased in HIV patient samples (Bhardawj et al., 2014; Contreras-Galindo et al., 2006). Recently, while this study was being undertaken, a subset of HERVs were shown to be upregulated in the Broncho alveolar lavage fluid (BALF) and peripheral blood mononuclear cells (PBMCs) of COVID-19 (coronavirus induced disease-19) patients (Balestrieri et al., 2021; Kitsou et al., 2021).

SARS-CoV-2 is a positive-strand RNA virus that causes COVID-19. The disease has caused numerous deaths and continue to be the top global public health concern. Within two years of its discovery, several virus variants have emerged. Some of them, such as the delta and the omicron variants are serious public health concerns (Tao et al., 2021; Skarbinski et al., 2022). Although multiple vaccines have been developed against SARS-CoV-2, there is limited access to vaccines globally and efficacy and duration of the protection offered by the vaccines vary from person to person. Antiviral therapeutics are the preferred treatment option in unvaccinated or vaccine-nonresponsive cases. It is important to understand the mechanisms of viral pathogenesis and decode the host-pathogen interactions to develop specific antiviral therapeutics against SARS-CoV-2.

Considering the importance of HERVs in inflammation and immune response pathways and the observed modulation of HERV expression in other viral infections, the current study was designed to systematically measure the expression of HERVs in SARS-CoV-2 infection-permissive human cell lines and PBMCs of COVID-19 patients. Among the different HERVs analyzed, only the HERV-R envelope level was significantly reduced in SARS-CoV-2 infected cells in all cases. Further studies confirmed an antiviral role of the HERV-R envelope in SARS-CoV-2 infected cells. Cross-talk between HERV-R and SARS-CoV-2 and its significance during SARS-CoV-2 infection is discussed.

## Results

### Profiling of HERV expression in human cell line-based models of SARS-CoV-2 infection

Earlier studies from our laboratory as well as other laboratories have demonstrated that Huh7 cells are permissive to SARS-CoV-2 infection (Verma et al., 2021). In order to monitor the effect of SARS-CoV-2 infection on the expression of HERVs, Huh7 cells were infected with the SARS-CoV-2 (NR-52281, SARS-related-coronavirus-2 isolate ISA-WA1/2020) at 0.1 MOI and infected cells were harvested after 72 hours and 96 hours post-infection. Mock infected cells were maintained in parallel as controls. RT-qPCR and western blot analysis of aliquots of the samples showed the presence of viral RNA and Nucleocapsid protein at both the time points, confirming infection of the cells (Fig. 1A, 1B). GAPDH RNA and protein levels were used as a reference for the normalization of the data (Fig. 1A,1B). RT-qPCR analysis of aliquots of the same samples using different HERV-specific primers revealed an increase in HERV-T, HERV-E, HERV-V, HERV-FRD, HERV-MER34, HERV-W and HERV-K-HML-2 envelope mRNAs at 72 hours post-infection (Fig. 1C, 1D, 1F, 1G, 1H, 1I, 1J). There was an increase in HERV-T, HERV-MER34, and HERV-K-HML-2 envelope mRNAs at 96 hours post-infection (Fig., 1C, 1H, and 1J). In contrast, there was a significant decrease in the HERV-R envelope mRNA level in SARS-CoV-2 infected cells at both 72- and 96-hour time points (Fig. 1E).

**Figure 1:**
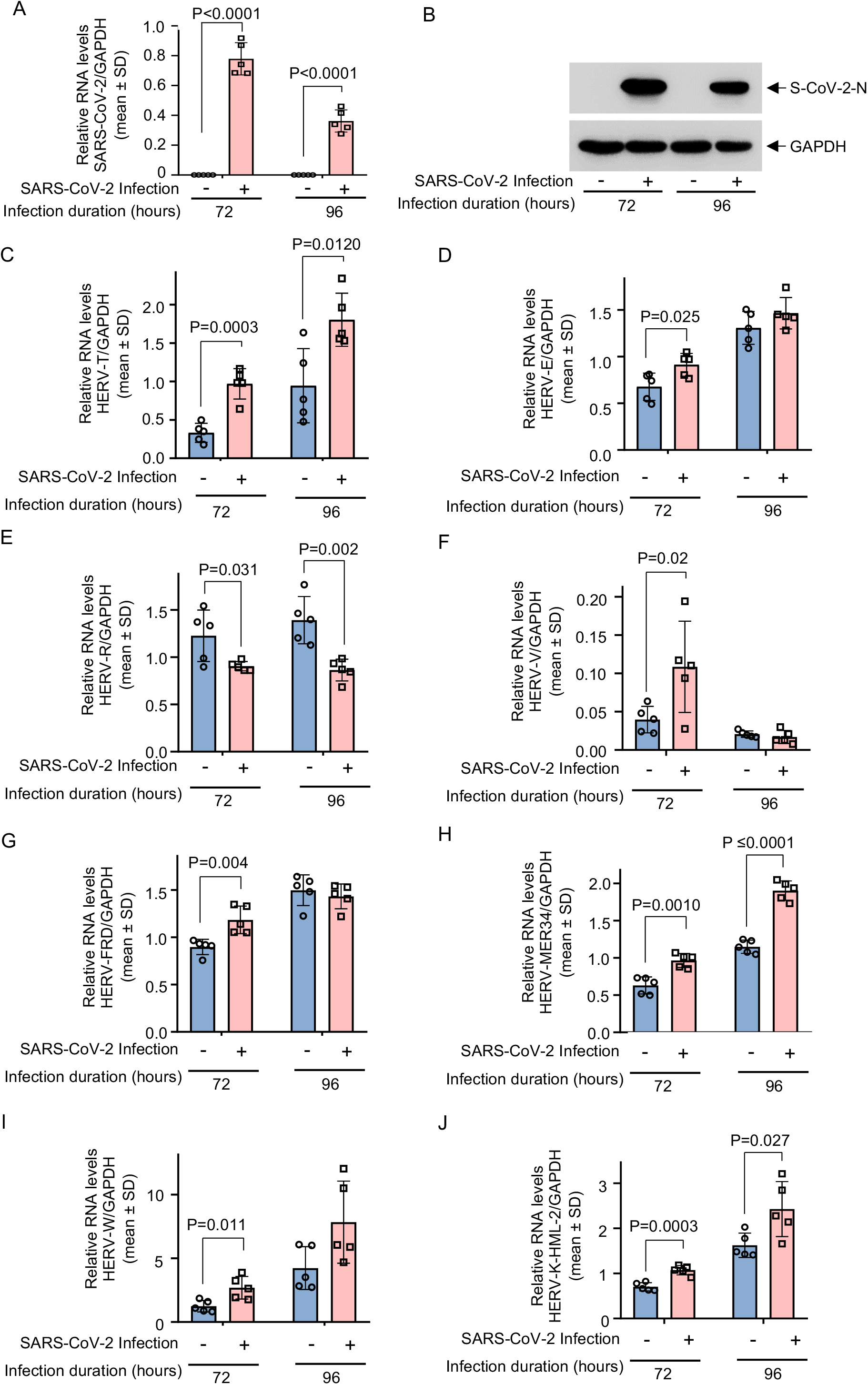
Analysis of HERV expression in the SARS-CoV-2 infected Huh7 cells. (A) RT-qPCR measurement of intracellular level of the SARS-CoV-2 RNA (normalized to that of the GAPDH) in Huh7 cells after 72 hours and 96 hours of infection. (B) Western blot analysis of the SARS-CoV-2 N protein (upper panel) and GAPDH (lower panel) levels in Huh7 cells, as mentioned in (A). (C-J) RT-qPCR analysis of the intracellular mRNA levels of HERV-T (C), HERV-E (D), HERV-R (E), HERV-V (F), HERV-FRD (G), HERV-MER34 (H), HERV-W (I) and HERV–K-HML-2 (J) in the SARS-CoV-2 infected Huh7 cells, as mentioned in (A). Values were normalized to that of the GAPDH and represented as mean±SD of samples from five independent experiments. Student’s t-test was applied to determine the significance between the groups.

Next, the pattern of HERV expression was evaluated in two other Human cell line-based models of SARS-CoV-2 infection. A human lung epithelial cell line (A549) and an acute monocytic leukemia cell line (THP-1) was selected. Since both A549 and THP-1 cells lack detectable level of ACE2 protein, replication-deficient Adenovirus encoding human ACE2 (hACE2) was transduced into these cells to make them permissive for SARS-CoV-2 infection. As expected, hACE2 mRNA and protein were detected in the Adeno-hACE2 transduced cells, respectively (Fig. 2A-2C). Twenty-four hours post-transduction, these cells were infected with the SARS-CoV-2 and incubated for additional 72 hours, followed by RT-qPCR and western blot mediated detection of the viral RNA and Nucleocapsid protein in these cells. As expected, a significant amount of viral RNA was detected only in the samples transduced with ACE2, followed by infection with the SARS-CoV-2 (Fig. 2D, 2E). Productive infection in hACE2 transduced cells was further confirmed by the detection of viral N-Protein in both A549 and THP-1 cells (Fig. 2F). RT-qPCR analysis of different HERVs in aliquots of the above samples showed a significant decrease in HERV-R envelope mRNA level in SARS-CoV-2 infected A549 cells (Fig. 2G). There was an increase in the mRNA level of HERV-E, V, FRD, MER34, W and K whereas HERV-T envelope mRNA remains unaltered (Fig. 2G). In the case of THP-1 cells, the HERV-R envelope mRNA level was significantly decreased and there was an increase in the mRNA level of the rest of the HERVs tested (Fig. 2H).

**Figure 2:**
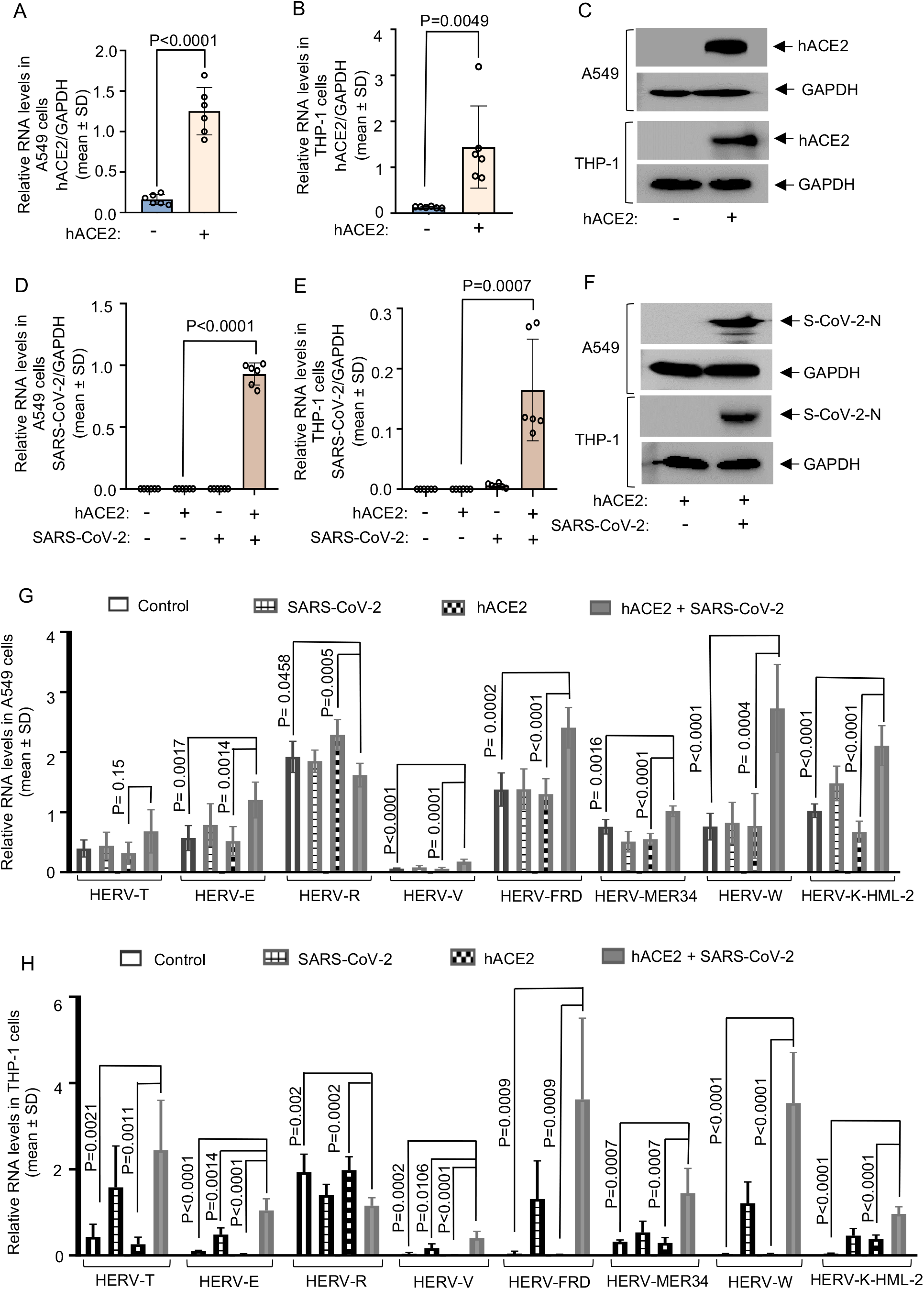
Analysis of HERV expression in the SARS-CoV-2 infected A549 and THP-1 cells. (A) RT-qPCR measurement of hACE2 mRNA level (normalized to that of the GAPDH) in the A549 cells transduced with hACE2 expressing adenovirus. (B) RT-qPCR measurement of hACE2 mRNA level (normalized to that of the GAPDH) in the THP-1 cells transduced with hACE2 expressing adenovirus. (C) Western blot analysis of the ACE2 (1^st^ and 3^rd^ panels) and GAPDH (2^nd^ and 4^th^ panels) protein levels in the hACE2 adenovirus transduced A549 and THP-1 cells, as indicated. (D) RT-qPCR measurement of intracellular level of the SARS- CoV-2 RNA (normalized to that of the GAPDH) in A549 cells transduced with hACE2 expressing adenovirus and infected with the SARS-CoV-2 for 72 hours, as indicated. (E) RT- qPCR measurement of intracellular level of the SARS-CoV-2 RNA (normalized to that of the GAPDH) in THP-1 cells transduced with hACE2 expressing adenovirus and infected with the SARS-CoV-2 for 72 hours, as indicated. (F) Western blot analysis of the SARS-CoV-2 N (1^st^ and 3^rd^ panels) and GAPDH (2^nd^ and 4^th^ panels) protein levels in A549 and THP-1 cells transduced with hACE2 expressing adenovirus and infected with SARS-CoV-2 for 72 hours, as indicated. (G) RT-qPCR measurement of various HERV mRNAs in A549 cells transduced with hACE2 expressing adenovirus, followed by infection with SARS-CoV-2 for 72 hours, as indicated. Values were normalized to that of the GAPDH and represented as mean±SD of 3 independent experiments. (H) RT-qPCR measurement of various HERV mRNAs in THP-1 cells transduced with hACE2 expressing adenovirus followed by infection with SARS-CoV-2 for 72 hours, as indicated. Values were normalized to that of the GAPDH and represented as mean±SD of 3 independent experiments. Student’s t-test was applied to determine the significance between the groups. P<0.05 was considered significant.

### SARS-CoV-2 infection inhibits the expression of HERV-R envelope protein

In order to evaluate the effect of SARS-CoV-2 infection on the protein level of the HERV-R envelope, Huh7 cells were infected with the SARS-CoV-2, the whole cell extract was prepared at 72- and 96-hours post-infection and western blotted using the anti-HERV-R envelope and anti-GAPDH antibodies. In agreement with RT-qPCR data, there was a significant reduction in HERV-R envelope protein level in the infected cells at both the time points (Fig. 3A). A more profound reduction in the HERV-R envelope level was observed in A549 and THP-1 cells upon infection with the SARS-CoV-2 at 72-hours post-infection (Fig. 3B).

**Figure 3:**
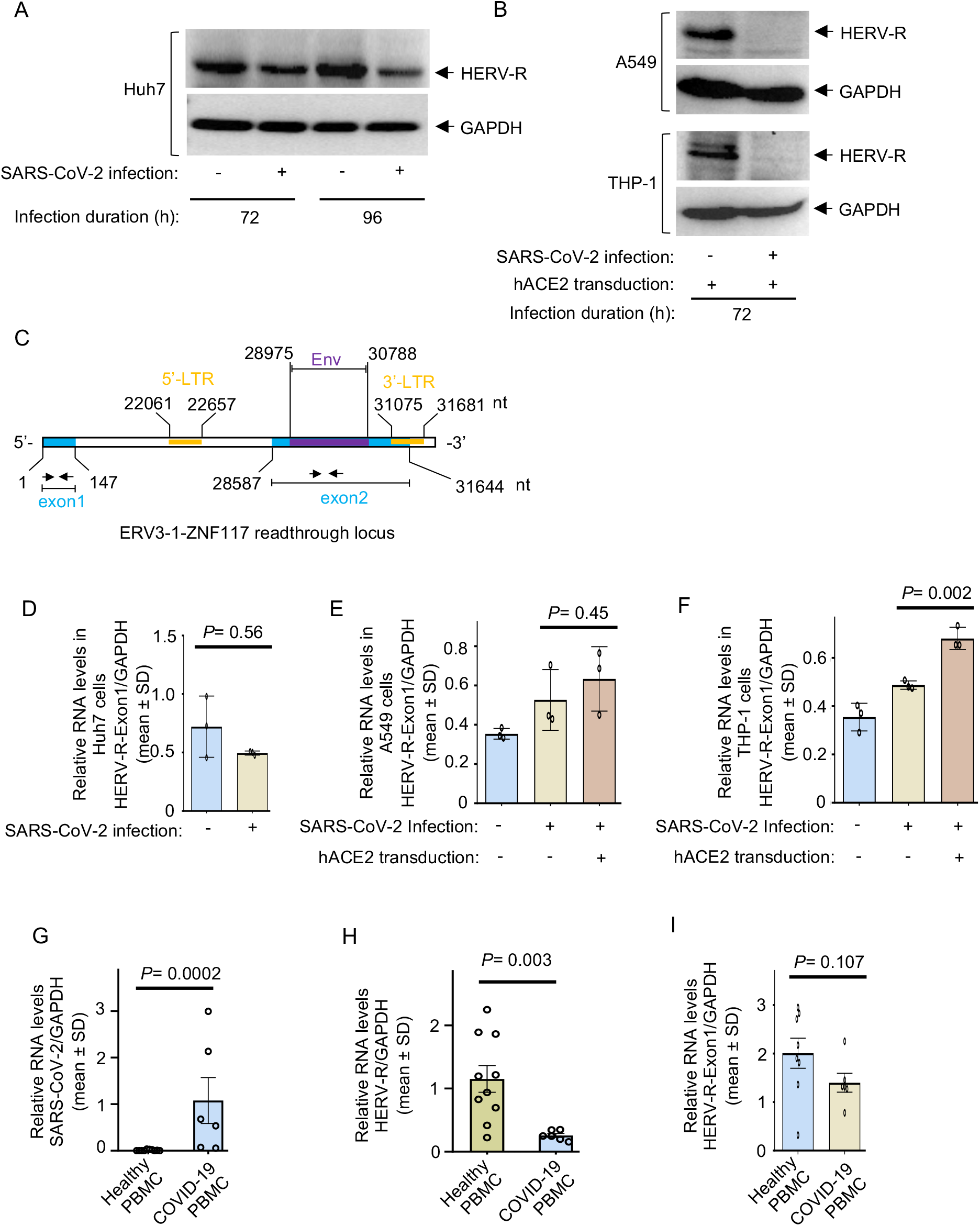
HERV-R envelope expression is decreased in SARS-CoV-2 infected cell lines and COVID-19 patient PBMCs. (A) Western blot analysis of the HERV-R envelope (upper panel) and GAPDH (lower panel) protein levels in Huh7 cells infected with SARS-CoV-2 for 72 and 96 hours, as indicated. (B) Western blot analysis of the HERV-R envelope (1^st^ and 3^rd^ panels) and GAPDH (2^nd^ and 4^th^ panels) protein levels in A549 and THP-1 cells transduced with hACE2 and infected with SARS-CoV-2 for 72 hours, as indicated. (C) Scheme of ERV3-1-ZNF117 readthrough genomic locus. Exon 1 and 2 belongs to ZNF117 and 5’-LTR, envelope (Env) and 3’-LTR belongs to HERV-R (ERV3-1). Numbers denote genomic coordinates of respective regions.(D) RT-qPCR measurement of exon 1 region [shown in (C)] transcript in Huh7 cells infected with SARS-CoV-2 for 72hours. (E) RT-qPCR measurement of exon 1 region [shown in (C)] transcript in A549 cells transduced with hACE2 expressing adenovirus, followed by infection with SARS-CoV-2 for 72hours. (F) RT-qPCR measurement of exon 1 region [shown in (C)] transcript in THP-1 cells transduced with hACE2 expressing adenovirus, followed by infection with SARS-CoV-2 for 72hours. (G) RT-qPCR measurement of intracellular level of SARS-CoV-2 RNA in PBMCs isolated from 10 healthy individuals and 6 COVID-19 patients.(H) RT-qPCR measurement of HERV-R envelope transcript level in PBMCs isolated from 10 healthy individuals and 6 COVID-19 patients. (I) RT-qPCR measurement of Exon1 region [shown in (C)] transcript level in PBMCs isolated from 10 Healthy individuals and 6 COVID- 19 patients. All RT-qPCR values were normalized to that of the GAPDH and represented as mean±SD of 3 independent experiments. Student’s t-test was applied to determine the significance between the groups.

Human genome contains around 40 HERV-R-like elements (Kannan et al., 1991; Andersson et al., 2005). However, only the HERV-R element located on chromosome 7 at 7q11, upstream of the zinc finger protein 117 (ZNF 117) locus, contains the complete open reading frame (ORF) for the viral envelope protein (Kannan et al., 1991). Other ORFs in this locus are nonfunctional due to nonsense mutations (Kannan et al., 1991). The ORF encoding the envelope protein (part of exon2) of HERV-R is flanked by Long-terminal repeats (LTRs) at both 5’- and 3’- ends (Fig. 3C). Read through transcription from HERV-R to ZNF117 have been indicated, suggesting the presence of an alternate promoter in the HERV-R locus (Bustamante Rivera et al., 2018). Exon 1 is located upstream of the HERV-R-5’-LTR, which may be referred to as a marker to distinguish between the activities of the HERV-R-LTR and the alternate promoter (Fig. 3C). RT-qPCR analysis of the HERV-R exon1 mRNA level in Huh7 cells revealed no change in the mRNA level upon infection with the SARS-CoV-2 (Fig. 3D). Similar results were obtained in the A549 cells (Fig. 3E). There was a marginal yet significant increase in the HERV-R exon1 mRNA level upon infection of THP-1 cells with the SARS-CoV-2 (Fig. 3F). Above data indicated that reduction in HERV-R envelope protein level in the SARS-CoV-2 infected cells may be attributed to reduced activity of the HERV-R LTR. We next measured the level of HERV-R envelope mRNA in PBMCs isolated from healthy individuals and COVID-19 patients. There was a significant decrease in the HERV-R envelope mRNA level in the PBMCs of COVID-19 patients whereas the exon-1 mRNA level was unaltered (Fig. 3G-3I). These data confirmed that the HERV-R envelope level is suppressed during the natural course of SARS-CoV-2 infection and further supported the role of HERV-R-LTR in controlling the expression of the HERV-R envelope.

### Overexpression of HERV-R envelope inhibits SARS-CoV-2 replication by activating the extracellular signal-regulated kinase

In order to test if HERV-R envelope overexpression modulates SARS-CoV-2 replication, Huh7 cells overexpressing the HERV-R envelope protein were infected with the SARS-CoV- 2 for 48 hours, followed by measurement of viral RNA and protein levels. There was a significant decrease in the viral RNA and nucleocapsid protein level in HERV-R envelope expressing cells (Fig. 4A, 4B). HERV-R has been reported to be involved in multiple cellular processes and pathways (Bustamante Rivera et al., 2018). To identify the mechanism underlying its antiviral activity against SARS-CoV-2, the effect of HERV-R overexpression on major cellular signaling pathways was checked. HERV-R overexpression did not affect antiviral or proinflammatory signaling pathways in HEK293T cells, measured by luciferase reporter assays (Fig. 4C). While constitutively active RIG-I mutant, R-C (harboring RIG-I CARD domains alone) induced production of firefly luciferase driven by IFNβ, ISRE, ISG56 and NFκB promoters, expression of HERV-R failed to induce any of these promoters (Fig.4C).

**Figure 4:**
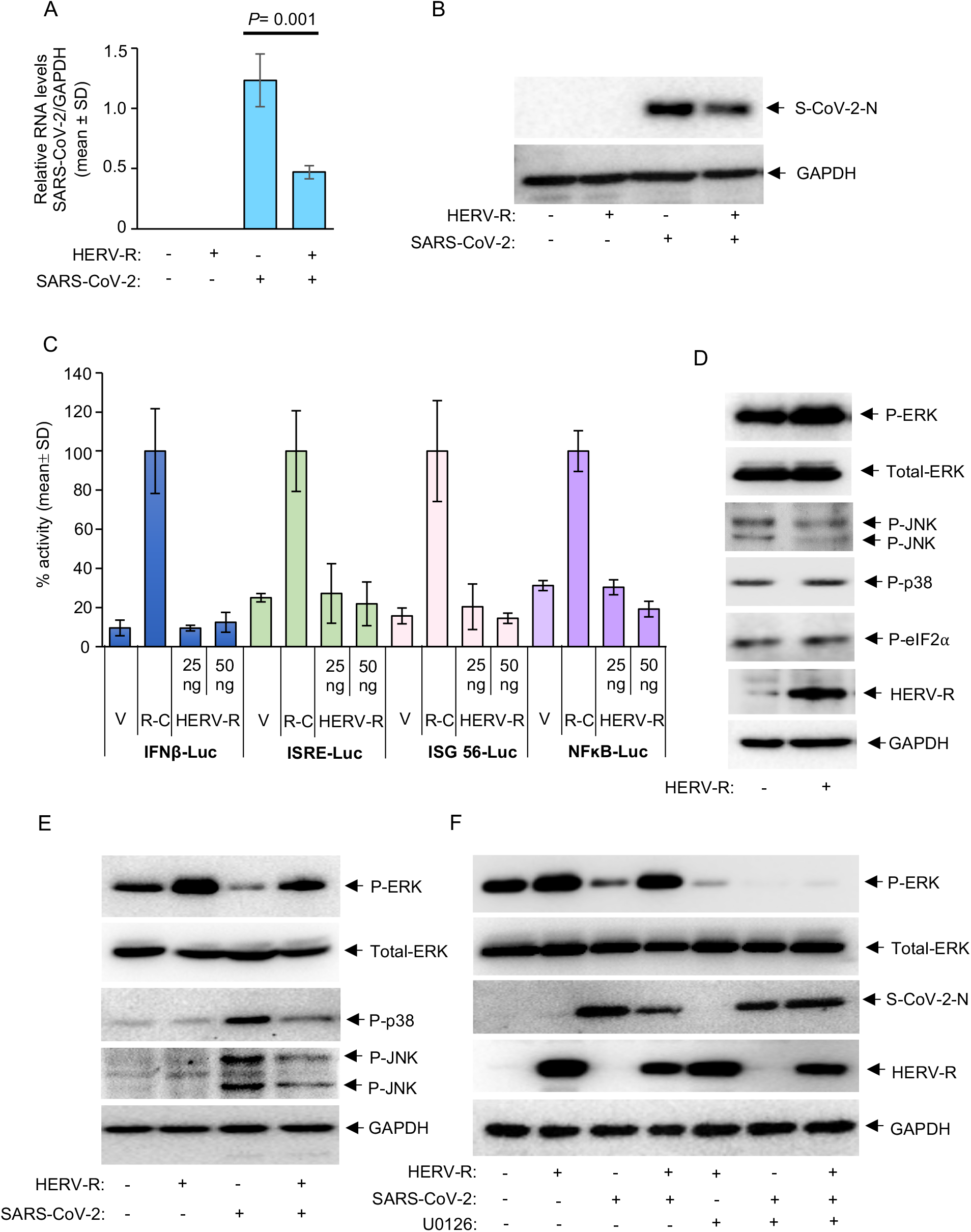
Activation of the extracellular signal-regulated kinase (ERK) by HERV-R envelope leads to inhibition of SARS-CoV-2 replication. (A) RT-qPCR measurement of intracellular level of the SARS-CoV-2 RNA (normalized to that of the GAPDH) in Huh7 cells transiently transfected with HERV-R expression plasmid and infected with the SARS-CoV-2. Values are mean±SD of 3 independent experiments. (B) Western blot analysis of the SARS-CoV-2 N protein (upper panel) and GAPDH (lower panel) levels in Huh7 cells, transiently transfected with HERV-R expression plasmid and infected with the SARS-CoV-2. (C) Dual luciferase assay showing activation of IFNγ, ISRE, ISG56 and NFκB promoters in HEK 293T cells expressing the empty vector (V), constitutively active mutant of RIG-I, R-C, 25 and 50 ng of HERV-R. Activation of these promoters by R-C was taken as 100%. Data are represented as mean percentage of FF Luc/RL Luc of triplicate samples ± SD. (D) Western blot analysis of the indicated proteins in Huh7 cells, transiently transfected with HERV-R expression plasmid. (E) Western blot analysis of the indicated proteins in Huh7 cells, transiently transfected with HERV-R expression plasmid and infected with the SARS-CoV-2. (F) Western blot analysis of the indicated proteins in Huh7 cells, transiently transfected with HERV-R expression plasmid, infected with the SARS-CoV-2 and treated with U0126.

Among the mitogen-activated protein kinase (MAPK) signaling pathway effectors, HERV-R activated the extracellular signal-regulated kinase (ERK), measured using phospho-ERK (Thr 202, Tyr 204) specific antibody (Fig. 4D, 1^st^ panel, denoted as P-ERK). Total-ERK level was checked as control (Fig. 4D, 2^nd^ panel). HERV-R overexpression did not alter the activity of c-jun N-terminal kinases (JNK-1-3) and p38 MAPK, two major stress-induced MAPKs (Fig. 4D, 3^rd^ and 4^th^ panel, denoted as P-JNK and P-p38, respectively). The level of phospho-eIF2α (Ser 51), which is the central mediator of the integrated stress response pathways, was also unaltered in HERV-R overexpressing Huh7 cells (Fig. 4D, 5^th^ panel, denoted as P-eIF2α). Level of HERV-R and GAPDH was measured as controls (Fig. 4D, 6^th^ and 7^th^ panel).

SARS-CoV-2 infection is known to upregulate the p38 MAPK pathway and inhibition of the p38 MAPK pathway blocks SARS-CoV-2 replication in infected cells (Bouhaddou et al., 2020). ERK, p38 and JNK MAPK pathways are known to antagonize the activity of each other depending on the upstream signal and cellular condition (Fey et al., 2012; Xiao et al., 2002; Junttila et al., 2008; Fey et al., 2012). Therefore, the cross-talk between ERK, p38 MAPK and JNK pathways was investigated in SARS-CoV-2 infected Huh7 cells overexpressing HERV-R. HERV-R overexpressing cells were infected with the SARS-CoV-2 and the levels of phospho-ERK, phospho-JNK and phospho-p38 MAPK was checked. There was a decrease in phospho-ERK (P-ERK) level in SARS-CoV-2 infected cells, which could be partially reversed in HERV-R overexpressing cells (Fig. 4E, 1^st^ panel). Total-ERK level was checked as control (Fig. 4E, 2^nd^ panel). The phospho-p38 MAPK (P-p38) level was upregulated in SARS-CoV-2 infected cells, which was not seen in cells overexpressing HERV-R (Fig. 4E, 3^rd^ panel). Interestingly, phospho-JNK (P-JNK) level was also significantly upregulated in SARS-CoV-2 infected cells (Fig. 4E, 4^th^ panel). There was a decrease in P-JNK level in SARS-CoV-2 infected cells overexpressing HERV-R (Fig. 4E, 4^th^ panel).

GAPDH level was checked as a control to ensure equal loading (Fig. 4E, 5^th^ panel). In order to confirm the role of ERK in mediating the antiviral function of HERV-R, Huh7 cells overexpressing HERV-R were treated with the MEK1/2 (mitogen-activated protein kinase kinase 1) inhibitor U0126, followed by measurement of P-ERK and viral Nucleocapsid protein level. As expected, the U0126 treatment significantly reduced the level of P-ERK in all samples (Fig. 4F, 1^st^ panel). Total-ERK level was checked as control (Fig. 4F, 2^nd^ panel). Viral nucleocapsid protein level was decreased in cells overexpressing HERV-R and infected with SARS-CoV-2, which was abrogated by U0126 treatment (Fig. 4F, 3^rd^ panel). HERV-R envelope and GAPDH levels were checked to ensure HERV-R expression and equal loading of samples (Fig. 4F, 4^th^ and 5^th^ panels). These data confirmed that antiviral effect of HERV-R against SARS-CoV-2 is mediated by its ability to induce the activity of ERK.

### Activation of p38 MAPK by the SARS-CoV-2 leads to inhibition of HERV-R envelope transcription

HERV-R envelope is flanked by LTRs at its 5’- and 3’- ends (Fig. 3C). The LTR is 596 nucleotides long and sequence of both 5’- and 3’- LTRs are identical. In order to test the functional activity of HERV-R LTR in mammalian cell lines, the LTR sequence (596 bp) was inserted upstream of the Gaussia-Luciferase (G-Luc) reporter coding sequence. SV40 Poly A was inserted downstream of the reporter sequence to ensure transcription termination and enhanced stability of the reporter mRNA (Fl 5’-LTR-Gluc, Fig. 5A). A vector lacking the LTR sequence was used as a control (G-Luc). A vector expressing the firefly-luciferase coding sequence under control of the SV40 promoter and SV40 Poly A sequence was used as an internal control for normalization of the G-Luc data. Significant G-Luc reporter activity was detected in Huh7 cells transfected with the Fl 5’-LTR G-Luc plasmid, supporting the role of HERV-R 5’-LTR in driving transcription (Fig. 5B). Next, a deletion mutant of LTR (22061- 22385 nucleotides deleted, Δ5’-LTR-G-Luc) was expressed in Huh7 cells in parallel with the Fl 5’-LTR-G-Luc reporter or G-Luc reporter. G-Luc activity of the Δ5’-LTR-G-Luc was reduced by more than 90% in Huh7 cells, when compared to that of the Fl 5’-LTR-G-Luc reporter, suggesting that indispensable transcription regulatory elements are located between 22061-22385 nucleotides of the HERV-R-LTR (Fig. 5C). Next, infection of Huh7 cells expressing the Fl-5’ LTR-G-Luc with SARS-CoV-2 showed ∼50% reduction in the reporter activity, in line with the reduction of HERV-R envelope RNA and protein level (Fig. 5D).

**Figure 5:**
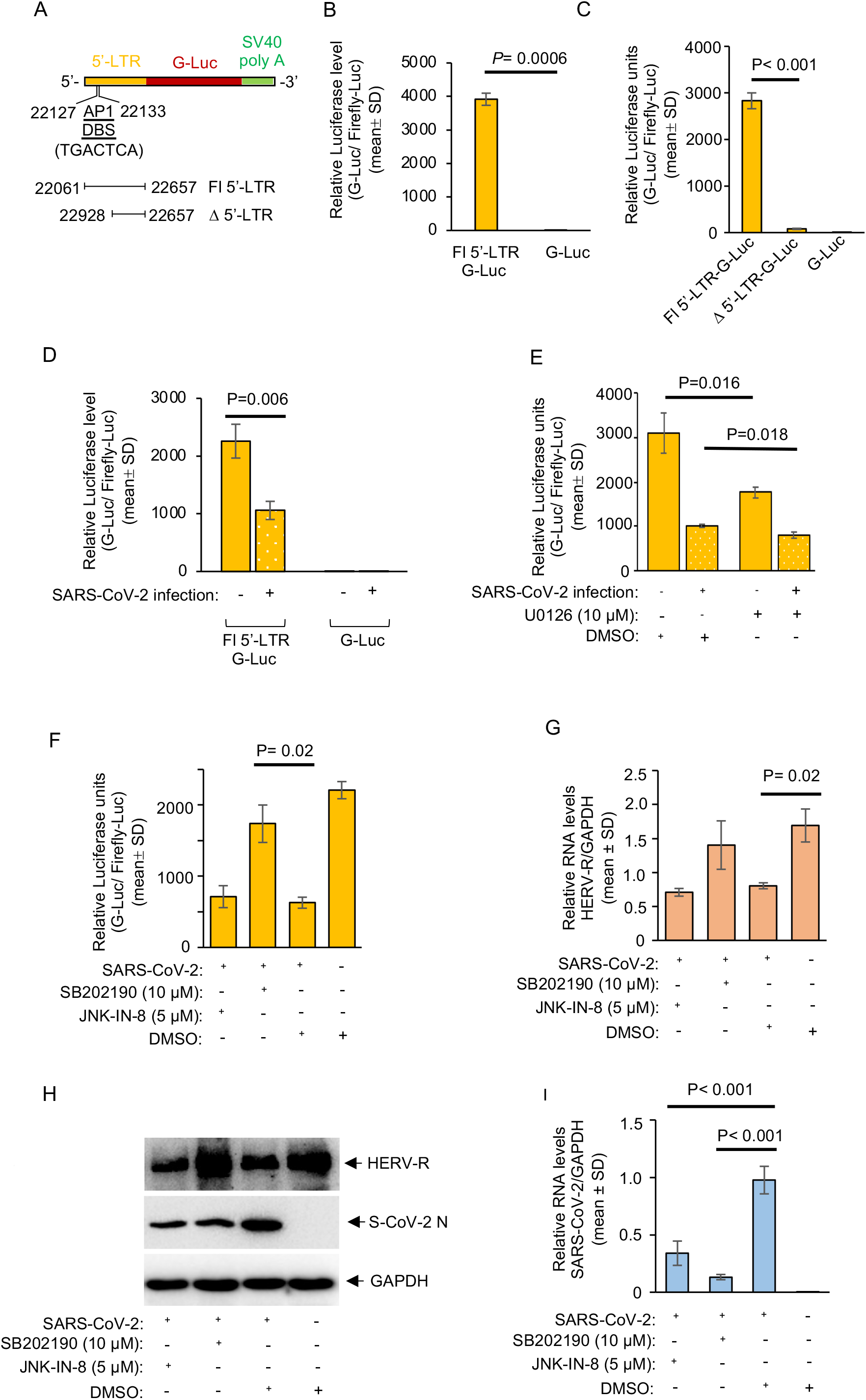
Expression of HERV-R envelope is inhibited by p38 MAPK activation and concomitant inhibition of ERK in SARS-CoV-2 infected cells. (A) Schematic of the Luciferase reporter cassette showing the position of full length 5’-LTR (Fl 5’-LTR, 22061-22657 nucleotides) and truncated 5’-LTR (Δ 5’-LTR, 22385-22657 nucleotides). Position and sequence of the AP1 DNA binding site present in the LTR is marked (AP1 DBS). (B) HERV-R 5’-LTR activity in Huh7 cells cotransfected with the pGL3 Fl 5’-LTR G-Luc and pGL3 promoter Firefly-Luc (denoted as Fl 5’-LTR G-Luc) or pGL3 G-Luc and pGL3 promoter Firefly-Luc (denoted as G-Luc) plasmids for 48 hours. The G-Luc values were divided by that of the Firefly-Luc and represented as mean (± SD) of three independent experiments. (C) HERV-R 5’-LTR activity in Huh7 cells cotransfected with the pGL3 Fl 5’- LTR G-Luc and pGL3 promoter Firefly-Luc (denoted as Fl 5’-LTR G-Luc) or pGL3 Δ 5’-LTR G-Luc and pGL3 promoter Firefly-Luc (denoted as Δ 5’-LTR G-Luc) or pGL3 G-Luc and pGL3 promoter Firefly-Luc (denoted as G-Luc) plasmids for 48 hours. The G-Luc values were divided by that of the Firefly-Luc and represented as mean (± SD) of three independent experiments. (D) HERV-R 5’-LTR activity in Huh7 cells cotransfected with the pGL3 Fl 5’- LTR G-Luc and pGL3 promoter Firefly-Luc (denoted as Fl 5’-LTR G-Luc) or pGL3 G-Luc and pGL3 promoter Firefly-Luc (denoted as G-Luc) plasmids, followed by infection with the SARS-CoV-2 for 48 hours. The G-Luc values were divided by that of the Firefly-Luc and represented as mean (± SD) of three independent experiments. (E) HERV-R 5’-LTR activity in Huh7 cells cotransfected with the pGL3 Fl 5’-LTR G-Luc and pGL3 promoter Firefly-Luc plasmids, followed by infection with the SARS-CoV-2 and treatment with U0126 for 24 hours. The G-Luc values were divided by that of the Firefly-Luc and represented as mean (± SD) of three independent experiments. (F) HERV-R 5’-LTR activity in Huh7 cells cotransfected with the pGL3 Fl 5’-LTR G-Luc and pGL3 promoter Firefly-Luc plasmids, followed by infection with the SARS-CoV-2 and treatment with the indicated compounds for 24 hours. The G-Luc values were divided by that of the Firefly-Luc and represented as mean (± SD) of three independent experiments. (G) RT-qPCR measurement of HERV-R envelope mRNA in Huh7 cells infected with the SARS-CoV-2 and treated with the indicated compounds for 48 hours. Values were normalized to that of the GAPDH and represented as mean±SD of 3 independent experiments. (H) Western blot analysis of the HERV-R envelope (upper panel), SARS-CoV-2 N (middle panel) and GAPDH (lower panel) protein levels in Huh7 cells infected with the SARS-CoV-2 and treated with the indicated compounds for 48 hours. (I) RT-qPCR measurement of intracellular level of the SARS-CoV-2 RNA (normalized to that of the GAPDH) in Huh7 cells infected with the SARS-CoV-2 and treated with the indicated compounds for 48 hours. Wherever shown, student’s t-test was applied to determine the significance between the groups.

Next, analysis of the HERV-R-LTR DNA sequence revealed the presence of several transcription regulatory elements corresponding to the DNA binding sites (DBS) of well-known transcription factors such as IRF1, IRF2, P53, E2F, Glucocorticoid receptor (GR) and CHOP etc. Importantly, a consensus AP1 DBS was also found between nucleotides 22127- 22133 (TGACTCA) in the HERV-LTR (Fig. 5A). AP1 family transcription factors such as c-Fos, c-Jun and ATF2 proteins are known to bind to the AP1 DBS and regulate transcription of the target genes. Different MAPKs activate subsets of AP1 proteins by phosphorylating them. For example, growth factor induced ERK-mediated phosphorylation of c-Fos and c-jun activates transcription of many immediate-early genes, including c-Fos and c-Jun (Garrington and Johnson, 1999; Karin et al., 1997). Similarly, stress-induced p38 MAPK/JNK mediated phosphorylation of ATF2 either activates or inhibits transcription of target genes, depending on the state of the cell (Zarubin and Han, 2005; Rong et al., 2010). Thus, AP1 DBS acts as a regulatory element for both transcription activation and repression.

In order to test the role of ERK, p38 MAPK and JNK in regulating the activity of HERV-R- LTR, Fl 5’-LTR-GLuc expressing Huh7 cells were infected with the SARS-CoV-2, followed by treatment with biochemical inhibitors of the MEK1/2 (U0126), p38 MAPK (SB202190) and JNK (JNK-IN-8). Treatment with U0126 significantly reduced HERV-R-LTR activity in Huh7 cells (Fig. 5E). SARS-CoV-2 infection inhibited the HERV-R-LTR activity, which was further decreased upon U0126 treatment (Fig. 5E). In contrast, treatment with SB202190 restored G-Luc activity in the infected cells, suggesting a role of p38 MAPK in inhibiting the HERV-R- LTR activity (Fig. 5F).

RT-qPCR and western blot analysis of the endogenous HERV-R RNA and protein levels in the presence of SB202190 in SARS-CoV-2 infected cells showed reversal of inhibition of HERV-R RNA and protein levels whereas JNK-IN-8 had no effect (Fig 5G, 5H).

Measurement of SARS-CoV-2 Nucleocapsid protein and RNA levels in aliquots of the above-mentioned samples showed a significant reduction in both protein and RNA levels (Fig. 5H, 5I). Taken together, these data demonstrate an important role of the ERK and p38 MAPK in mediating the antiviral effect of the HERV-R envelope protein.

## Discussion

An in-depth understanding of the host-pathogen cross-talk is important for designing specific prophylactic and therapeutic strategies against the pathogen. Here, we focused on exploring the cross-talk between HERVs and the SARS-CoV-2. Expression analysis of a few representative members of class I and class II HERVs was carried out in three different cell-based models of SARS-CoV-2 infection and PBMCs isolated from COVID-19 patients. Two epithelial cell derived lines such as Huh7 and A549 cells lines and a monocyte derived cell line (THP-1) was used for the infection studies. Of the eight HERVs tested, the RNA level of seven HERVs were upregulated in SARS-CoV-2 infected Huh7 and THP-1 cells (HERV-T, HERV-E, HERV-V, HERV-FRD, HERV-MER34, HERV-W, HERV-K-HML-2) whereas no change in HERV-T level was seen in SARS-CoV-2 infected A549 cells. The level of the HERV-R envelope was decreased in all three cell-based infection models as well as in the PBMCs isolated from COVID-19 patients. While this study was being undertaken, two publications reported the modulation of a subset of HERVs in COVID-19 patients. Balestrieri et al. reported upregulation of HERV-W envelope RNA and protein levels in T-lymphocytes of COVID-19 patients (Balestrieri et al., 2021). In another study, Kitsou et al. analyzed the RNA seq data of bronchoalveolar lavage fluid (BALF) and PBMC of COVID-19 patients and compared that to healthy individuals. HERV-FRD, HERV-H, HERV-W, ERV-L, HERV-I and HERV-K (HML-5, HML-3, HML-1) were significantly upregulated in the BALF of COVID-19 patients. No difference was seen in the level of HERV-E, HERV-K (HML-2, HML-4, HML-6) and HERV-9 (Kitsou et al., 2021). In the case of PBMCs, significant downregulation was seen in ERV-L, HERV-FRD, HERV-H and HERV-I whereas no change was seen in the level of HERV-K (HML1-6), HERV-W, HERV-9 and HERV-E (Kitsou et al., 2021). Expression profiles of HERV-W, HERV-FRD and HERV-K-HML-2 were upregulated in our cell based models of SARS-CoV-2 infection as well as in the BALF samples of COVID-19 patients reported by (Kitsou et al., 2021). In addition, we observed upregulation of HERV-E, HERV-V, HERV-MER34 and downregulation of HERV-R envelope in all three cell types. Our data suggest that alteration in HERV profiles is directly linked to SARS-CoV-2 infection rather than an indirect outcome of COVID-19 disease manifestation. Furthermore, our study shows that the cell-based infection models of SARS-CoV-2 are useful for investigating the cross- talk between SARS-CoV-2 and HERVs.

Since only the HERV-R envelope was downregulated in all three cell-based models and COVID-19 patient PBMCs, we investigated the significance of the HERV-R envelope in the life cycle of SARS-CoV-2. Interestingly, overexpression of the HERV-R envelope showed an inhibitory effect on SARS-CoV-2 replication. HERV-R/ERV3 is a class I ERV, related to gamma-retroviruses. It is found in Hominidae (excluding Gorilla) and Cercopithecoidea and is located at an identical genomic position (Chromosome 7q11) in great apes, monkeys and humans (Bustamante Rivera et al., 2018). HERV-R is expressed in many tissues including lung epithelium, spleen, lymph nodes, thymus, stomach, intestine, placenta etc. HERV-R has been reported to stimulate the immune system, trigger inflammatory pathways and involved in autoimmunity. It is also upregulated in many cancers (Bustamante Rivera et al, 2018).

We observed that HERV-R expression does not activate antiviral or proinflammatory signaling in the Huh7 cells under normal cellular conditions. However, it activates the ERK pathway, which is the major prosurvival MAPK pathway. Our data clearly shows that the cognate viral LTR drives the transcription of HERV-R envelope. We also observed the presence of a consensus AP1 DBS in the HERV-R-LTR and deletion of this region from the LTR abrogated its activity. Since HERV-R activates the ERK pathway, which controls the synthesis and activation of AP1 transcription factors such as c-Fos and c-Jun, we speculate that AP1 DBS is the key cis-acting element that drives the synthesis of HERV-R envelope through a positive feedback loop.

Among the HERVs, the envelope protein of HERV-K-HML2 has been shown to activate the ERK pathway through its cytoplasmic tail (Lemaitre et al., 2017). Further, an immunosuppressive peptide derived from the envelope protein of retroviruses (CKS-17) has been shown to activate the ERK pathway (Takahashi et al., 2001). HERV-K-HML2 envelope was shown to act upstream of Raf kinase to activate the ERK pathway (Lemaitre et al., 2017). CKS-17 was shown to act upstream of MEK to activate the ERK pathway (Takahashi et al., 2001). HERV-R envelope protein contains an N-terminal heptad repeat, a CKS17-like immunosuppressive region, a CX6C motif, and a C-terminal heptad repeat. It is possible that HERV-R CKS17-like immunosuppressive region is responsible for the activation of the ERK pathway. Since the ERK pathway plays a key role in both immune signaling and cancer, a clear understanding of the mechanistic details of ERK activation by the HERV-R envelope will provide critical insight into its biological role.

Earlier studies have shown the activation of the p38 MAPK pathway in SARS-CoV-2 infected cells. Our results are in agreement with the previous finding. In addition, our study shows that SARS-CoV-2 infection activates the JNK pathway and treatment with JNK inhibitor blocks viral replication equally effectively as the p38 MAPK inhibitor. Hence, a combination of p38 MAPK inhibitor and JNK inhibitor might be a more effective antiviral against the SARS-CoV-2. Although both p38 MAPK and JNK inhibitors block SARS-CoV-2 replication, only p38 MAPK inhibitor restores HERV-R envelope expression. Hence, HERV-R-LTR activity seems to be controlled by the cross-talk between ERK and p38 MAPK pathways.

The antagonism between ERK and p38 MAPK pathways has been reported earlier. Activation of ERK promotes cell survival and proliferation and inhibits apoptosis. On the other hand, activation of p38 MAPK inhibits ERK activity via protein phosphatase 1 or protein phosphatase 2A, thereby reversing the effect of ERK (Xiao et al., 2002; Junttila et al., 2008). Also, JNK and p38 MAPK mediated ATF2 activation has been shown to repress transcription of target genes by recruiting Histone deacetylase 4 (HDAC 4) to the promoter region (Rong et al., 2010). Thus, one possibility is that in SARS-CoV-2 infected cells, the HERV-R envelope is not transcribed due to a block in its LTR activity owing to a lack of ERK activity, which limits the synthesis of AP1 transcription factors. The other possibility is that increased p38 MAPK activity in SARS-CoV-2 infected cells activates the ATF2 transcription factor, which binds to the AP1 DBS located in the HERV-R-LTR and prevents HERV-R envelope transcription by recruiting transcription repressors. The above possibilities might coexist to minimize HERV-R envelope expression in SARS-CoV-2 infected cells. Future studies should clarify the actual mechanism.

In summary, the current study demonstrates the utility of human cell line-based SARS-CoV- 2 infection models in elucidating the cross-talk between HERVs and SARS-CoV-2 and evaluates the expression profile of a subset of HERVs in the above models. Importantly, our study also reveals the role of the HERV-R envelope protein as a host restriction factor against the SARS-CoV-2. These findings illustrate yet another evolutionary benefit of the integration of retroviral elements into the human genome.

## Methods

### Plasmids and reagents

5’-LTR of HERV-R was cloned in fusion with the downstream Gaussia Luciferase (G-Luc) into the pGL3 basic vector (Promega, USA). The sequence corresponding to 596 nucleotides of HERV-R 5’-LTR (22061-22657 base pairs on human chromosome 7q11) was PCR amplified from the genomic DNA of Huh7 cells. G-Luc coding sequence was PCR amplified from the G-Luc expression plasmid. Fusion cloning was carried out by overlap PCR. The primers used for 5’-LTR are: FP- 5’ TACAAGAAGCTTTATATGAGGCAGGAAATATAAAAGG 3’; RP- 5’ ACCCACCATGGTCAGAAAAGGCTTGACAGTAAACCTGTAGTCTCC 3’ and for G-Luc, FP- 5’ TCAAGCCTTTTCTGACCATGGTGGGTATGAACATGCTACTCATCATGGGAGTCAAAGTT CTGTT 3’; RP- 5’ TTTCCTGCTTCATTCCCCCCTTTTTAGTCACCACCGGCCCCCTTG 3’. These two PCR products were fused by overlap PCR and cloned into the pGL3 basic vector between HindIII and XbaI sites (denoted as Fl 5’-LTR-G-Luc). Δ 5’-LTR-G-Luc clone was generated by restriction digestion of the Fl 5’-LTR-G-Luc with AflII (blunted) and SmaI, gel extraction of the 4kb fragment and self-ligation. All clones were confirmed by DNA sequencing. pGL3 promoter vector was from Promega (USA).

Mammalian expression plasmid encoding the untagged HERV-R envelope sequence was procured from Sino Biological (Cat no. HG19174-UT, Sino Biological Inc, China). IFNβ-Luc, ISRE-Luc, ISG56-Luc and NFκB-Luc, pRL-CMV and R-C has been described earlier (Hingane et al., 2020). Anti-P-p38 MAPK (Cat No. 4511), anti-P-JNK (Cat No. 58328), anti-P-ERK (Cat No. 9101) and anti-total-ERK (Cat No. 4695) antibodies were from Cell signalling Technology (Massachusetts, USA). Anti-GAPDH antibody (Cat No. SC-25778) was from Santacruz Biotechnology (Texas, USA). Anti-P-eIF2α antibody (Cat no. STJ92521) was from St John’s Laboratory, (London, UK). Anti-SARS-CoV-2-N antibody was from GeneTex Inc (USA). Goat anti-rabbit IgG-HRP (Cat No. 4030-05) and Goat anti-mouse IgG-HRP (Cat No. 1030-05) was from Southern Biotech (Alabama, USA). U0126 (Cat no. V112A) was from Promega (Wisconsin, USA). SB202190 (Cat no. S7067) was from Sigma (Missouri, USA). JNK-IN-8 (Cat no.S4901) was from Sellekchem (Texas, USA).

### Mammalian cell culture

Huh7 cells were cultured as described earlier (Nair et al., 2016). They were originally obtained from the laboratory of Prof Charles Rice (Blight et al., 2000). A549 and THP-1 cells were obtained from ATCC, USA. Huh7 and A549 cells were maintained in Dulbecco’s modified Eagle medium (DMEM) while THP-1 cells were cultured in RPMI-1640 medium. The media was supplemented with 10% FBS and Penicillin, Streptomycin during maintenance.

### PBMC isolation

Blood Samples were collected as per the recommended guidelines of the Institutional Ethics Committee of THSTI and ESIC (Employees state insurance corporation) Hospital, Faridabad (IEC Ref No: THS 1.8.1/ (97) dated July 07, 2020). Venous blood was drawn from six symptomatic COVID-19 patients (∼0-3 days from PCR positive report) and ten age-matched healthy participants at ESIC Hospital, Faridabad. The peripheral blood mononuclear cells (PBMCs) were isolated as described previously (Binayke et al., 2022). Briefly, blood was collected in sodium heparin-coated CPTTM tubes (BD Biosciences, USA) after receiving written informed consent from the donors. The tubes were centrifuged at 1500g for 25 min and the PBMCs were separated, washed twice with PBS, and used for RNA isolation, as described below.

### Adenovirus transduction and SARS-CoV-2 infection

Human ACE2 was expressed in A549 and THP-1 monocytes by adenovirus transduction, following published protocol (Hassan et al., 2020). Replication deficient adenovirus expressing human ACE2 (Ad5CMVhACE2, 2*10^11^ pfu/mL stock, denoted as hACE2 hereafter) were obtained from the Viral Vector Core (University of Iowa, Iowa, USA). 0.5*10^6^ A549 or 10^6^ THP-1 cells in serum free media were transduced with 10 MOI of hACE2 along with 8µg/ml polybrene for 1 hour at 37^0^C, in a 5% CO_2_ incubator, followed by washing of cells with PBS and SARS-CoV-2 infection. No cytotoxicity was observed in hACE2 transduced cells. SARS-CoV-2 was obtained from BEI Resources (NR-52281, SARS-related coronavirus-2 isolate ISA-WA1/2020), amplified in Vero E6 cells in the BSL3 facility of THSTI, India, titrated and stored frozen in aliquots. SARS-CoV-2 infection was done as described (Verma et al., 2021). For hACE2 transduced cells, SARS-CoV-2 infection was done 24 hours post-transduction. In case of HERV-R over expression study, Huh7 cells were transfected with respective DNA 24 hours prior to infection with SARS-CoV-2. In case of inhibitor treatment, 10µM (final concentration) U0126, 10µM SB 202190 or 5µM JNK-In-8 was added to the culture medium after removing the infection medium and maintained for 48 hours, followed by collection of culture medium and/or cells for subsequent experiments. In inhibitor treated samples, 24 hours after addition of inhibitor, culture medium was replaced with fresh medium and inhibitor and incubated for further 24 hours.

### RNA isolation, Real Time-quantitative PCR (RT-qPCR), Western blot analysis

Total RNA was isolated using TRI reagent (MRC, Massachusetts, USA), followed by reverse transcription (RT) using Firescript cDNA synthesis kit (Solis Biodyne, Estonia). The relative transcript levels of HERVs or other genes were determined by SYBR green based RTq-PCR as described earlier (Verma et al., 2021). Values obtained for GAPDH RNA was used as an internal control for normalization of the data. Primers for quantification of SARS-CoV-2 are as described (Verma et al., 2021). Primers for quantification of HERVs are as described (Vincendau et al., 2015). Some of the primers were designed in this study using primer design tool available in Snapgene (Snapgene, USA) and Primer Blast. Primer sequences are: SCoV2 FP: 5’-TGGACCCCAAAATCAGCGAA 3’, SCoV2 RP: 5’- TCGTCTGGTAGCTCTTCGGT 3’; HERV-T FP: 5’ CCCCTACCCTTTTTGGGG 3’, HERV-T RP: 5’ GTACCCCAGGTAGGAAACTCTGGG 3’ ; HERV-E FP: 5’ GCTTTCTTTCTGATCCTAGGCTGTG 3’, HERV-E RP: 5’ CTTTGGGGAGGCGTTGGCTCGAGACC 3’ ; HERV-R FP: 5’ ATGTCGGGTCAAAGGAAGG 3’, HERV-R RP: 5’ GAATCGGTGGAACAAGCAG 3’; HERV-V FP: 5’ CCCCTCCTGGCTATGTATTT 3’, HERV-V RP: 5’ GCTCTTTTCTGTCTGGGTTGG 3’; HERV-FRD FP: 5’ TCTCATTCTCACGCCTTCAC 3’, HERV-FRD RP: 5’ CGCCTCTATGCTTGTCCATT 3’; HERV-MER34 FP: 5’ TAAATGGTCTGGGCGATGTG 3’, HERV-MER34 RP: 5’ GGTGGATTGTCTGTGTCTCCT 3’; HERV-W FP: 5’ TGAGTCAATTCTCATACCTG 3’, HERV-W RP: 5’ AGTTAAGAGTTCTTGGGTGG 3’; HERV-K-HML-2 FP: 5’ GGCCATCAGAGTCTAAACCACG 3’, HERV-K-HML-2 RP: 5’ CTGACTTTCTGGGGGTGGCC 3’; ACE2 FP: 5’ TGTAACTGCTGCTCAGTCC 3’, ACE2 RP: 5’ CCCATTTTGCTGAAGAGCC 3’; HERV-R Exon 1 FP: 5’ AGCCGGAGCTTCTGGTGTAGT 3’, HERV-R Exon 1 RP: 5’ CATTTCTAGGCTTCCAGTGGGTC 3’; hGAPDH FP: 5’ GAGTCAACGGATTTGGTCGT 3’, hGAPDH RP: 5’ TTGATTTTGGAGGGATCTCG 3’.

For western blot analysis, cells were harvested in Laemlli buffer (2% SDS, 13% glycerol, 2.5% 2-mercaptoethanol, 0.005% bromo phenol blue and 63 mM Tris HCl, pH ∼ 6.8) and incubated at 95^0^C for 10 minutes. Total protein amount was estimated by Bicinchoninic acid assay followed by resolution of equal amount of protein by SDS-PAGE and transfer to Polyvinylidene fluoride (PVDF) membrane. Blocking and incubation with different primary and secondary antibodies were done as per the guidelines provided by the manufacturer of the primary antibody. Bands were developed by enhanced chemiluminescence reagent, using a commercially available kit (BioRad, USA) and visualized in a gel Documentation system (Chemidoc MP, BioRad, USA).

### Dual Luciferase reporter assay

Huh7 cells were seeded at 70% confluency overnight and transfected with the Fl 5’- LTR G-Luc or Δ 5’-LTR G-Luc and pGL3 promoter-Firefly-Luc plasmids using lipofectamine 2000 at 1:1 (w/v) ratio. Transfection media was changed after 6 hours and cells were incubated for 48 hours. In case of inhibitor treatment, media was changed after 30 hours of transfection and indicated amount of compounds were added for 18 hours. In case of SARS-CoV-2 infection experiment, 24 hours post-transfection, cells were infected with 0.1 MOI of SARS-CoV-2 stock as described in the relevant section and incubated for 48 hours. For measurement of G-Luc activity, culture media was collected and subjected to measurement of Luciferase activity using the Renilla-Luciferase assay kit, following manufacturer’s protocol (Promega, Wisconsin, USA). For measurement of Firefly-Luc activity, cells were lysed, followed by measurement of Luciferase activity using the Firefly-Luciferase assay kit, following manufacturer’s protocol (Promega, Wisconsin, USA).The G-Luc values were divided by that of the Firefly-Luc and graphs were plotted as mean (± SD) of three independent experiments done in triplicates.

For IFN-β, ISRE, ISG56 and NFκB promoter activation, cell-based luciferase assay was performed in HEK293T cells in white-walled 96-well plates as reported earlier (Madhvi et al., 2017). Briefly, HERV-R plasmid (25 and 50 ng) was transfected into HEK293T cells along with reporter plasmids harboring Firefly luciferase driven by either i) interferon-β ii) ISRE iii) ISG56 or iv) NFκB promoter. In addition, as an internal control, a plasmid containing Renilla luciferase driven by a thymidine kinase (TK) promoter was also transfected. As a positive control for activation of these reporters, a constitutively active mutant of RIG-I, R-C (only the CARD domains) was used. All transfections were performed using Lipofectamine 2000 following manufacturer’s protocol. Luciferase activity was measured 24 h post-transfection using Promega Dual Glo luciferase assay kit following manufacturer’s protocol using Synergy HT Multi-Mode microplate reader (Bio-Tek, USA). Ratio of firefly to Renilla luciferase values was converted to percentages and the data was plotted as % activity. Values obtained for R-C, was considered as 100% and rest of the values were normalized to R-C values.

### Statistical analysis

Data are represented as mean±SD of three or more independent experiments (as indicated in figures) done in triplicates. Student’s t-test was applied for the cell line-based data while Mann Whitney U test was used for data involving human samples. A *p* value <0.05 with 95% confidence interval was considered statistically significant.

## Acknowledgement

This study was funded by the THSTI core grant to MS. AA acknowledges the Mission COVID Suraksha grant (BT/CS0010/CS/02/20) from DBT BIRAC, India. NG acknowledges funding support (TRAIN fellowship) from THSTI. SA, RV and ONS gratefully acknowledge the Council of Scientific and Industrial Research, Govt of India for providing senior research fellowship. SARS-Related Coronavirus 2, isolate USA-WA1/2020 (NR-52281) was deposited by the Centers for Disease Control and Prevention and obtained through BEI Resources, NIAID, NIH, USA.

## Author contributions

Experimental design and data analysis were done by NG, SA and MS. NG, SA and MS wrote the manuscript. NG, SA, RV, ONS, MKY, AB and KJ performed experiments. AA, SM, S, BN and CTRK contributed reagents. All authors edited the manuscript.

